# DILI_*C*_: An AI based classifier to search for Drug-Induced Liver Injury literature

**DOI:** 10.1101/2022.02.12.480184

**Authors:** Sanjay Rathee, Meabh MacMahon, Anika Liu, Nicholas Katritsis, Gehad Youssef, Woochang Hwang, Lilly Wollman, Namshik Han

**Affiliations:** Milner Therapeutics Institute, University of Cambridge, Cambridge, UK; LifeArc, Stevenage, UK; Centre for Molecular Informatics, Department of Chemistry, University of Cambridge, UK; Cambridge Centre for AI in Medicine, University of Cambridge, Cambridge, UK

**Keywords:** DILI, Drug Induced Liver Injury, NLP, Natural language processing, ML, Machine learning, AI, Artificial Intelligence

## Abstract

Drug-Induced Liver Injury (DILI) is a class of Adverse Drug Reactions (ADR) which causes problems in both clinical and research settings. It is the most frequent cause of acute liver failure in the majority of western countries and is a major cause of attrition of novel drug candidates. Manual trawling of literature for is the main route of deriving information on DILI from research studies. This makes it an inefficient process prone to human error. Therefore, an automatized AI model capable of retrieving DILI-related papers from the huge ocean of literature could be invaluable for the drug discovery community. In this project, we built an artificial intelligence (AI) model combining the power of Natural Language Processing (NLP) and Machine Learning (ML) to address this problem. This model uses NLP to filter out meaningless text (e.g. stopwords) and uses customized functions to extract relevant keywords as singleton, pair, triplet and so on. These keywords are processed by apriori pattern mining algorithm to extract relevant patterns which are used to estimate initial weightings for a ML classifier. Along with pattern importance and frequency, an FDA-approved drug list mentioning DILI adds extra confidence in classification. The combined power of these methods build a DILI classifier (DILI_*C*_) with 94.91% cross-validation and 94.14% external validation accuracy. To make DILI_*C*_ as accessible as possible, including to researchers without coding experience, an R Shiny App capable of classifing single or multiple entries for DILI is developed to enhance ease of user experience and made available at https://researchmind.co.uk/diliclassifier/).

## 1 INTRODUCTION

Drug-Induced Liver Injury (DILI) is a class of adverse drug reactions (ADR) which is an issue in both clinical and research settings. Although DILI can be mild, resolving once administration of the problem drug is discontinued, it lies on a spectrum and can also be severe. DILI is the most frequent cause of acute liver failure in the majority of western countries Hoofnagle and Björnsson (2019) and is a major cause of attrition of novel drug candidates Church and Watkins (2018) and accounts for almost one quarter of clinical drug failures Watkins (2011). As new findings on DILI are often published in scientific literature, collating this data from literature is useful for risk-assessment during drug development and in the clinic. However, currently manual trawling of text from literature is the main route of obtaining relevant information about DILI from research studies. This is an inefficient process prone to human error and modern computational techniques for mining textual data can improve it, a model capable of retrieving DILI-related papers from the huge ocean of literature could be invaluable for the drug discovery community.

Natural language processing (NLP) involves using computational techniques to extract information and insights from text data. Previous studies have applied NLP techniques to identify relevant literature for challenges in drug discovery, including with the goal of drug repurposing Zhu et al. (2020) and collating information on COVID-19 for researchers Wang and Lo (2021). Additionally previous attempts have been made to classify adverse drug events using NLP on available data Harpaz et al. (2014). Databases of drug side effects also contain DILI-related informationFDA (2021); Kuhn et al. (2016). In this study NLP is used to extract relevant patterns from literature and this knowledge is combined with information related to DILI from publicly available databases. This combined information is used to train a classifier to classify literature as DILI-related or not. Figure 1 highlights the flow of text processing for our model.

**Figure 1.**
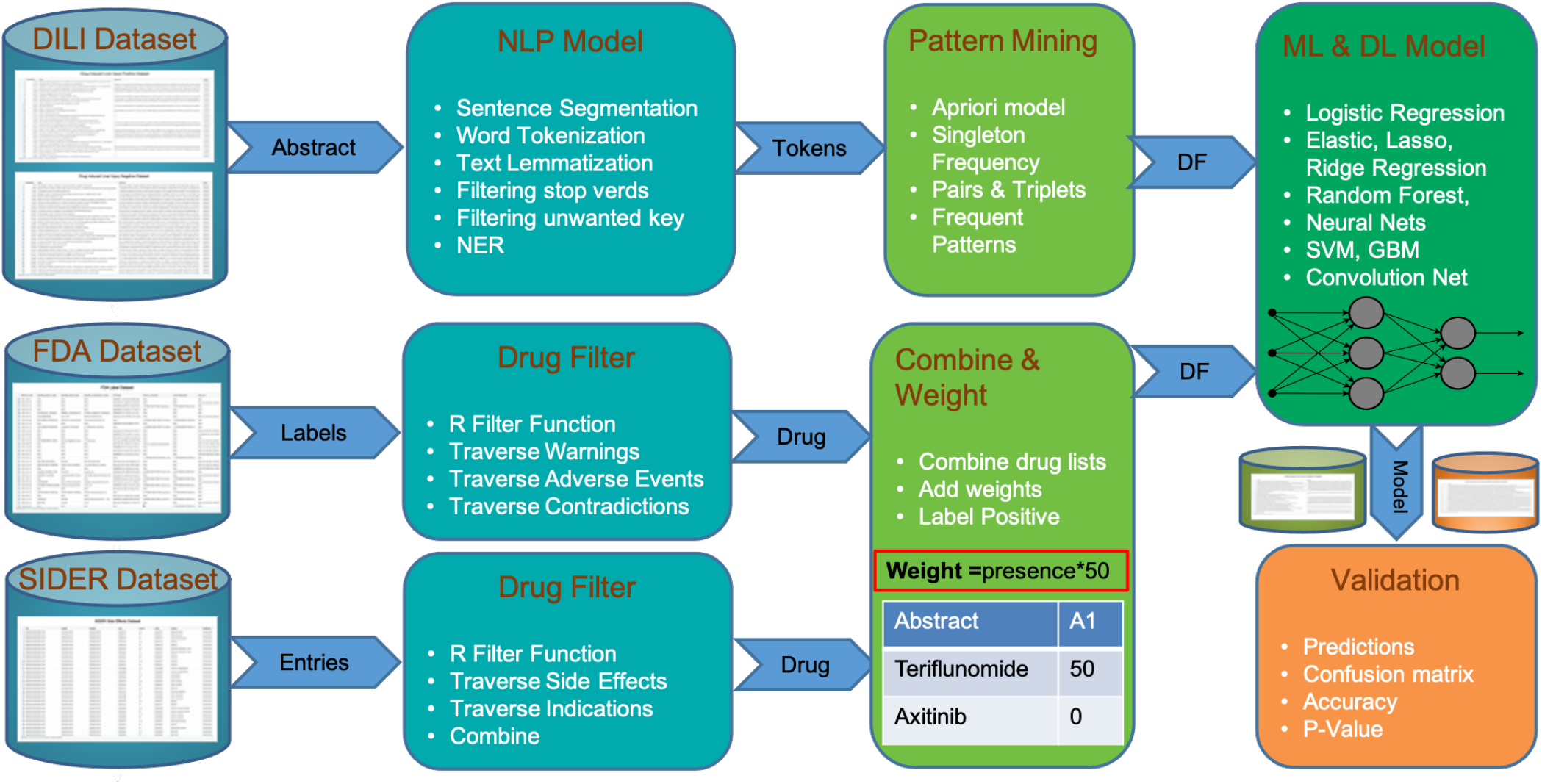
The steps of DILI_*C*_ from dataset of DILI positive and DILI negative papers to validations showing intergration of FDA and SIDER datasets.

## 2 MATERIALS AND METHODS

We built an artificial intelligence (AI) model combining the power of Natural Language Processing (NLP) and Machine Learning (ML) to extract relevant literature for DILI from ocean of published papers. This model combines the information available in the title and abstracts of scientific papers with information from external databases to improve the efficacy and accuracy. A detailed procedure is available in Algorithm 1 which contains all the steps to build this model.

### Algorithm 1 Classify Literature as DILI Positive or DILI Negative

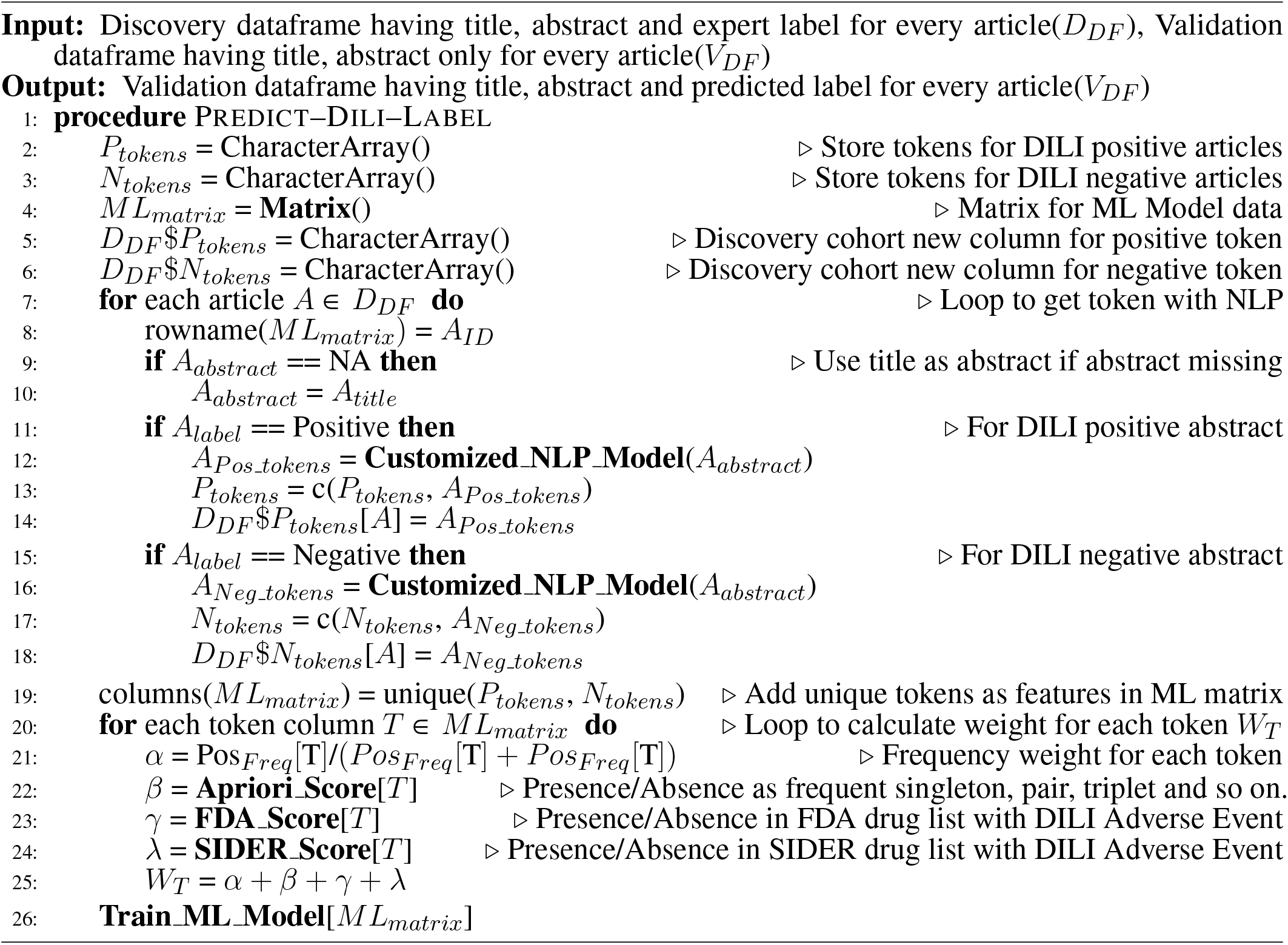

### 2.1 Data Preparation

A well curated dataset of ∼28,000 DILI annotated papers was obtained from the CAMDA team CAMDA (2021). This dataset was generated after filtering out the most obvious DILI literature which makes the task of classification challenging, but more representative of the challenge of sorting through real word literature beyond the obviously DILI related or entirely unrelated papers. All the papers in this dataset are labeled as DILI related (DILI positives) or not related to DILI (DILI negatives) by an experienced panel of experts. We used approximately half of this data with a balanced split of DILI positive and negative to extract insights and train a model (discovery set). The remaining half was kept as validation set.

We divided the discovery set of 14,203 papers into training (80%) and testing (20%) sets consistent with their labels. Overall, we used 5,741 DILI positive & 5,620 DILI negative as a training set and 1,436 DILI positive & 1,406 DILI negative as test set.

### 2.2 Natural Language Processing Model

A NLP model with some customization was used to extract the relevant information from the available training cohort (Algorithm 2). It starts with the most basic NLP step sentence tokenization on titles and abstracts, followed by word tokenization. A customized word tokenization method was developed to generate keyword sets of singleton, pairs, triplets and so on. This step generates combinations containing only nouns and adjectives and filters out irrelevant text like stop words using R UDPipe package. These keyword sets were processed for text lemmatization and stemming to generalise the list. The output of this NLP model was a vector containing all keyword sets as features and for each of these their frequency and length (singleton, pair) was stored as weights for pattern mining. This NLP model was applied on both title and abstracts.

#### Algorithm 2 Customized NLP Model to extract Tokens from Abstract

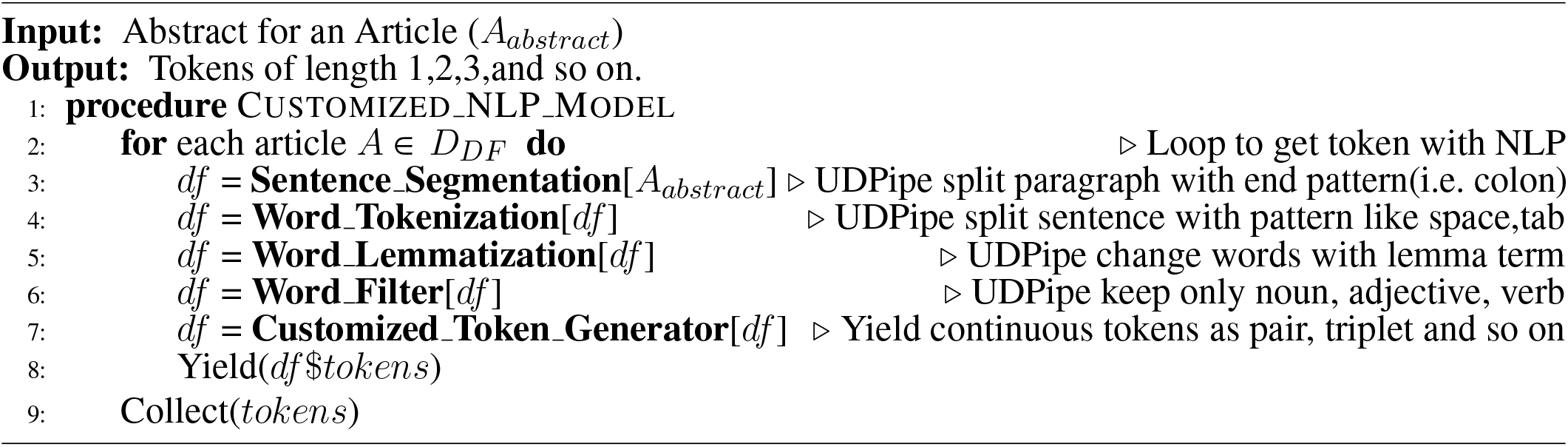

### 2.3 Pattern Mining

Along with the total frequency of a keyword set, the frequency of the keyword and its subsets in terms of the number of papers (DILI positive or DILI negative) in which it appears was calculated. The pattern mining ML algorithm Apriori was used for this. In this way, we included the frequency of a keyword set and its subset as a factor for weighting that keyword set. A distributed processing-based implementation of Apriori was used to minimize the overall processing time.

### 2.4 External Cohort Integration

Since external datasets contain information which could be advantageous in classifying DILI literature, two were integrated into the model. These two publically available datasets were the FDA approved drugs list FDA (2021) and SIDER adverse events dataset Kuhn et al. (2016). From these two datasets (Algorithm 3), a list of drugs with DILI as adverse events or warning were extracted, and these drugs were given a higher weight than others without such warnings. The side effects field of SIDER database for drugs was helpful to add extra information into this highly weighted list.

### 2.5 Classifier

The final vector of keywords along with their updated weights was given as an input to various well-known ML & AI models (Logistic Regression, Elastic Net, Random Forest, Neural Net, Support Vector Machine, Gradient Boosting Machine, Convolution Neural Networks and LSTM) to train a classifier. The weight of a keyword was calculated by its total frequency, length, FDA and SIDER list presence or absence.

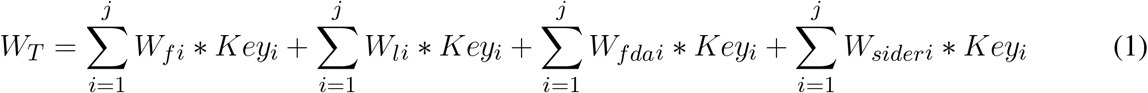

In equation (1), *W*_*T*_ represents the total weight for a paper, *key* represents the weight for presence(1) or absence(0) of a keyword set, (*W*_*f*_) represents the weight for frequency of a keyword set, *W*_*l*_ represents the weight for length of a keyword set (for instance singleton 1, pair 2, triplet 3), *W*_*fda*_ represents the weight for presence and absence in FDA list with DILI adverse event and *W*_*sider*_ represents the weight for presence and absence in SIDER list with DILI adverse event. The classifier with the highest cross-validation accuracy (Gradient Boosting Machines) was tested on a put-aside test set. The results on the test set were quite promising with an accuracy of 94.89%. The model was iterated 10 times with different test set to get the average accuracy of 94.9%. Figure 2 shows the probability of every sample being positive. Any sample with a probability higher than 50% is labelled as DILI positive. The cutoff of 50% can be adjusted to closer reflect a real-world dataset which will have far more negative literature. Table 1 shows the confusion matrix for the stand-out (20%) testing set.

**Figure 2.**
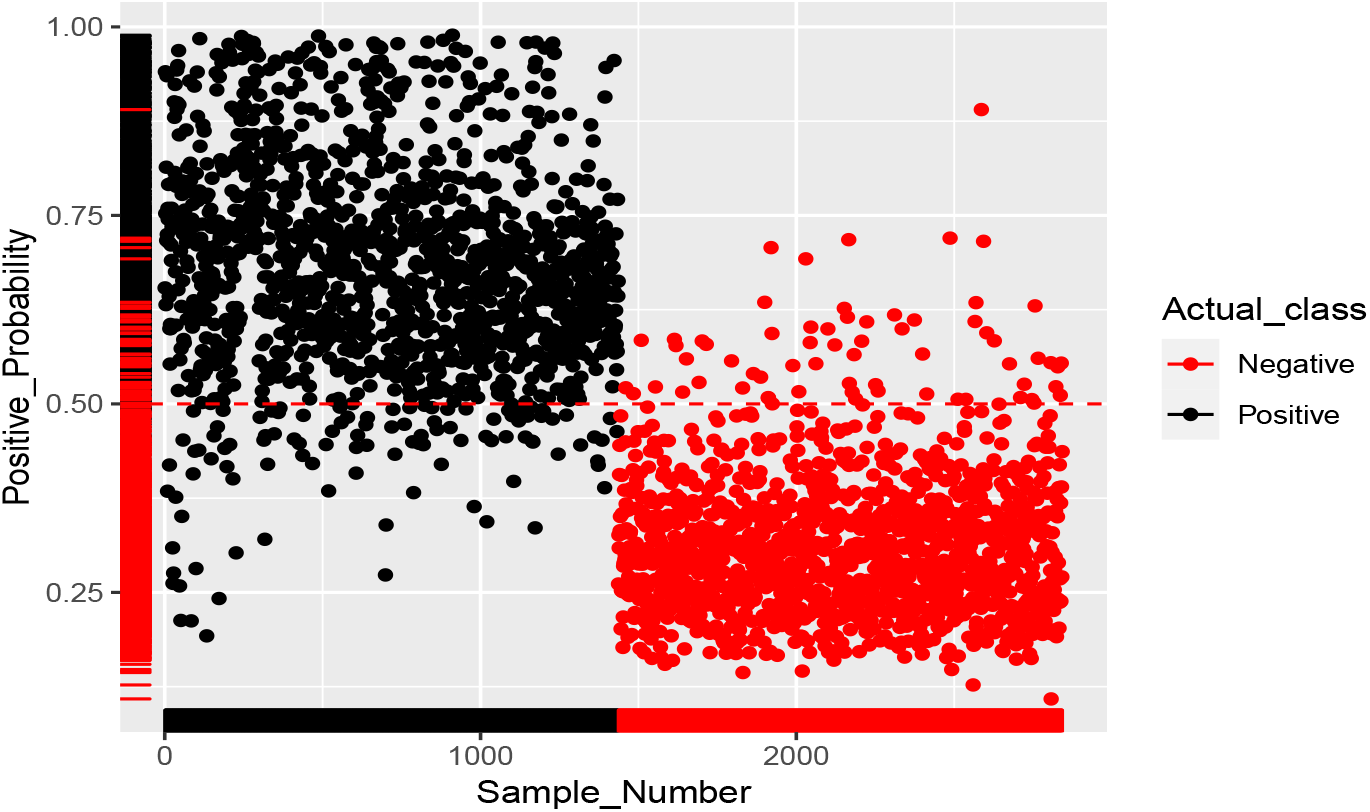
Prediction Probabilities Plot.

**Table 1.**
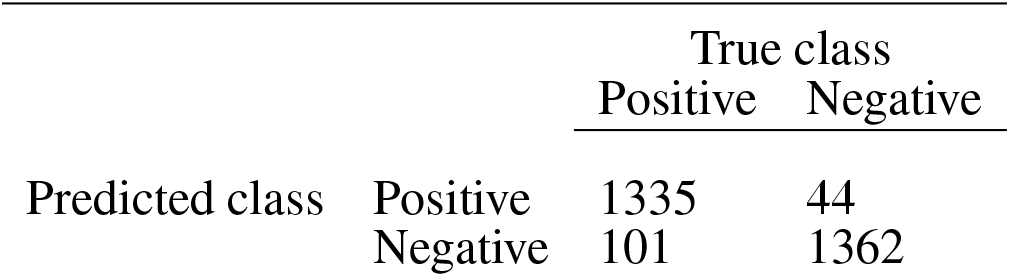
Confusion matrix of GBM classifier applied to stand out abstract cohort

|  |  | True class |  |
| --- | --- | --- | --- |
|  |  | Positive | Negative |
| Predicted class | Positive | 1335 | 44 |
|  | Negative | 101 | 1362 |

#### Algorithm 3 Add score for presence/absence in external cohort FDA and SIDER

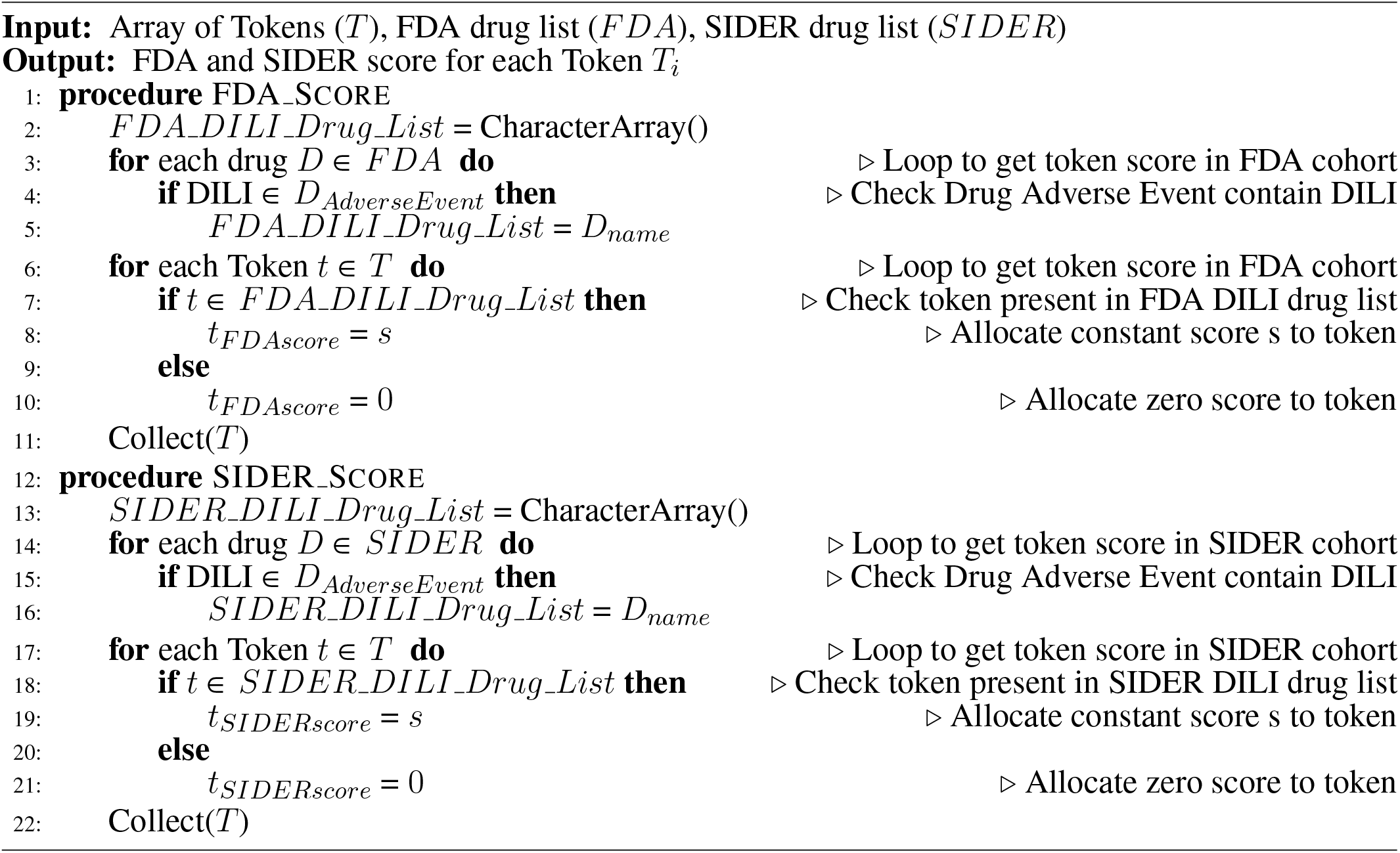

## 3 RESULTS

The most effective model was Gradient Boosting Machines (Figure 3) with 94.76% accuracy when applied to the internal hold out test set of 2,842 papers, half of which were DILI positive and half DILI negative. The inclusion of FDA and SIDER datasets improved the accuracy of the GBM model in the validation set and on an additional external set (Table 2). The final model is used to predict the labels for the external validation cohort shared by CAMDA. We got encouraging results with an accuracy of 94.14% and F1-Score 94.08%. The highlight of the model was its recall value of 96.02%. Dili_*C*_ was then applied to an unseen additional external set which was unbalanced DILI cohort, making it more reflective of real world data. On the additonal external set accuracy was 90.25% and an F1-score of 90.94%. The recall value was improved with this set, with a vaule of 97.9%

**Figure 3.**
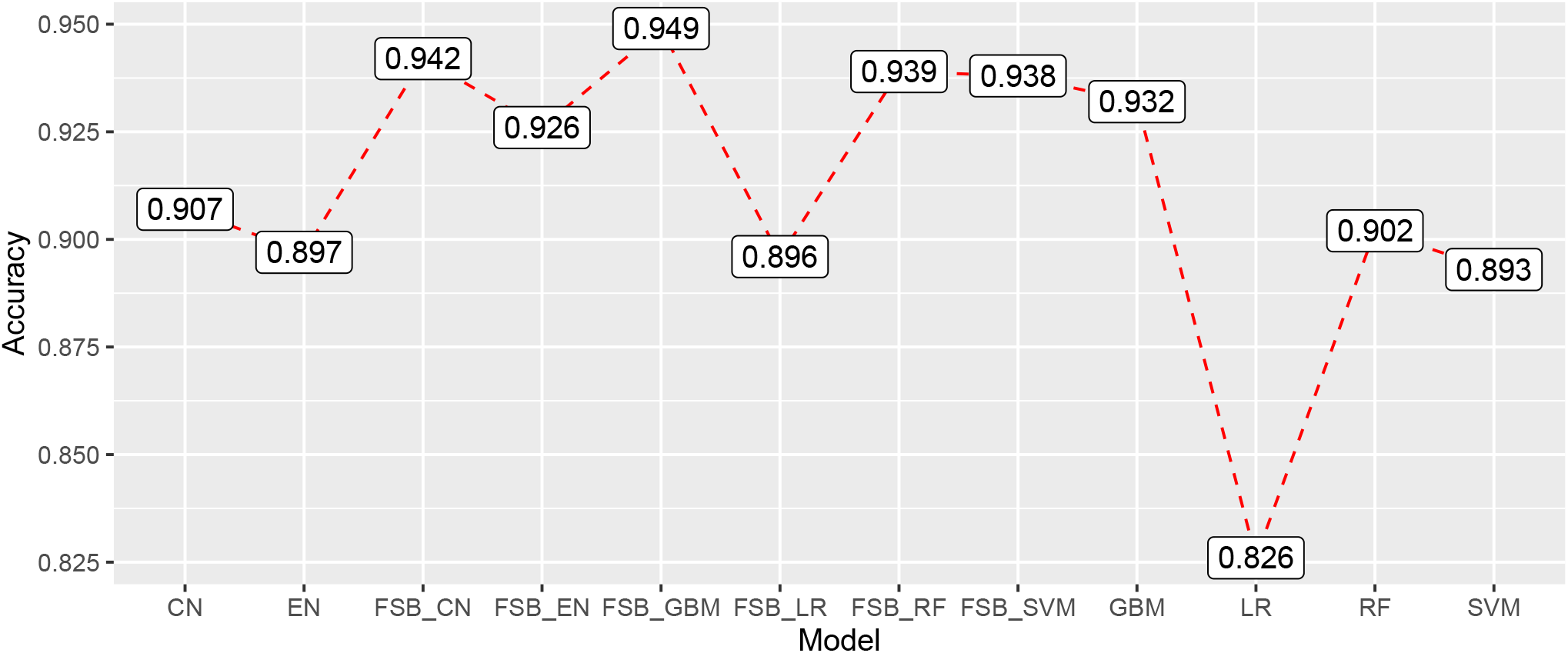
Internal accuracies for all ML classifiers (EN: Elastic Net, LR: Logistic Regression, SVM: Support Vector Machines, CN: Convolution Network, RF: Random Forest, GBM: Gradient Boosting Machines, FSB: Feature Selection Based Model) showing that GBM has the highest accuracy.

**Table 2.** Results for the GBM model applied to the validation set and additional external set of DILI and non-DILI literature. The inclusion of FDA and SIDER datasets improved the GBM model

|  | Validation Set (14211) |  |  |  | Additional external set (2000) |  |  |  |
| --- | --- | --- | --- | --- | --- | --- | --- | --- |
|  | Accuracy | F1 score | Recall | Precision | Accuracy | F1 score | Recall | Precision |
| GBM (abstract only) | 0.9386 | 0.9376 | 0.9631 | 0.9133 | 0.8845 | 0.8936 | 0.9700 | 0.8284 |
| GBM(+FDA) | 0.9406 | 0.9396 | 0.9659 | 0.9147 | 0.8915 | 0.8992 | 0.9680 | 0.8395 |
| GBM (+SIDER) | 0.9414 | 0.9408 | 0.9602 | 0.9221 | 0.9025 | 0.9094 | 0.9790 | 0.8491 |

## 4 DISCUSSION

DILI_*C*_ is a model with high accuracy which is useful to the community to classify literature as related to or unrelated to DILI, which can help do DILI risk-assessment for drugs during development, repurposing or in the clinic. Although it was developed to classify DILI literature, it has been designed to handle any adverse event classification problem so it’s has applications for drug risk-assessment beyond just liver injury to toxicities in other tissues. We note that complex machine learning AI models are known to have the power to magnify weak signals. In order to minimise the pressure on ML models and reduce the risk of such erroneous magnification, during the development of DILI_*C*_ a strong focus was put on the data cleaning processing steps of the model. Another potential issue is the chance that the inclusion of SIDER dataset could introduce bias against publications relating to drugs which aren’t yet included therein. Reassuringly, even without the inclusion of this database, DILI_*C*_ performs well, with an accuracy of 94.06% on the Validation set and of 89.15% on the additional external set. There is still potential to improve DILI_*C*_ in the future. Later steps like customized word segmentation, pattern mining, and external relevant cohorts add power to DILI_*C*_ and there is still plenty of scope to adjust the weights for these steps. In addition, as other databases related to drug toxicity and side effects are developed, these could be integrated to improve the model. To make DILI_*C*_ as accessible as possible, including to researchers without coding experience, an R Shiny App capable of classifing single or multiple abstracts for DILI is developed to enhance ease of user experience and made available at https://researchmind.co.uk/diliclassifier/).

## 5 CONCLUSIONS

DILI_*C*_ is a novel tool to classify literature as related to DILI or not. This is significant as it has the potential to aid researchers in drug-development and research settings during risk-assessment.

DILI_*C*_ is implemented in such a way that it can be modified to classify any other drug adverse reaction like DILI. Therefore, DILI_*C*_ code available at GitHub could be useful for researchers working in the same domain. A shiny app for DILI_*C*_ makes it user-friendly.

## CONFLICT OF INTEREST STATEMENT

SR is funded by JW Pharmaceutical. MM is an employee of Lifearc. NH and WH are funded by LifeArc. AL is funded by GlaxoSmithKline. NH is a cofounder of KURE.ai and CardiaTec Biosciences and an advisor at Biorelate, Promatix, Standigm and VeraVerse.

## AUTHOR CONTRIBUTIONS

SR, MM and NH conceived and designed the analysis; SR and MM collected the data, built the model, performed the analysis and wrote the paper; AL,NK, GY, WH and LW contributed data or analysis tools; AL and NH reviewed the paper; NH supervised the project.

## FUNDING

SR is funded by JW Pharmaceutical, MM and WH are funded by LifeArc, AL is funded by GlaxoSmithKline, NK is funded by Cambridge Trust

## ACKNOWLEDGMENTS

The authors thank CAMDA for provision of curated DILI datasets for training and testing.

## DATA AVAILABILITY STATEMENT

The code datasets analysed for this study can be found in the DILI Github. [https://github.com/sanjaysinghrathi/DILI-Classifier].

## Notes

### Competing Interest Statement

The authors have declared no competing interest.

